# Avifauna composition in Rice plantations: effects of Harvesting Stage and Surrounding Vegetation. Insights from Weppa Farm, Agenebode

**DOI:** 10.1101/2024.10.20.619302

**Authors:** Ilyas Ibrahim, Benhilda Antonio, Damen Tongken, Bethel Clement, Barnabas Sani Bello, Othniel Ugomeh, Jemima Amos Samuel, Hassanatu Patrik, Chukuma Dike

## Abstract

Rice plantations provide important habitats for diverse avian communities which support both resident and migratory species. These ecosystems serve as breeding grounds, provide foraging resources and temporary habitats for various bird species, including waterbirds, shorebirds, and passerines. This study was conducted in Weppa Farm (total farm size ∼13000 hector), Agenebode, Etsako East local government area in Edo State, Nigeria to investigate the avifauna compositions in pre-harvested and post-harvested rice plantations and the influence of surrounding vegetation on avian abundance. Four line transects were surveyed around the rice plantation. The transects were 400m long and split into 4, 100m sections. Surveys were done on the rice field before harvest (Unharvested), during harvest (partially harvested) and post-harvest (harvested). Surveys were done in the morning (7:00-9:00 GMT) and repeated in the evening (16:00-18:00 GMT). Vegetation parameters were measured at the beginning of each section of the transect. A total of 48 different bird species from 6 feeding guilds were recorded. Granivorous birds were the most abundant feeding guild (n=599). The Unharvested rice plantation had the highest abundance (n=1453), highest richness (r=37) but not diversity (H=5.18), partially harvested farm had the least abundance (n=592) and least diversity (H=4.95). The surrounding vegetations had a significant effect on bird abundance. Granivores mainly weavers were the most abundant species. The were mainly observed foraging and using weeds as nesting material. Insectivorous birds were the second must abundant species and could serve as biocontrol for arthropod invasions on the farm. Surrounding vegetation influences the abundance of birds in a rice plantation. There was high bird abundance in areas that had trees and shrubs nearby.

## INTRODUCTION

Rice (*Oryza glaberrima*) is extensively consumed worldwide, and the demand is growing in Africa. It is the most consumed grain in West Africa (Sottomayor et al., 2024;Adjah et al., 2022). Avian biodiversity in rice ecosystems differs significantly, with species richness varying by farming stage and season (Amira et al., 2018). Insectivores and granivores are the most common feeding guilds observed in most rice plantations (Amzah et al., 2021). Rice fields provide important habitats for diverse bird species and support both resident and migratory populations (Amira et al., 2018;Odoukpe et al., 2014). These ecosystems serve as breeding grounds, provide foraging resources, and temporary habitats for various bird species, including waterbirds, shorebirds, and passerines (Pierluissi, 2010). While some birds, like queleas, may be considered pests, many species provide ecosystem services such as pest control (Amzah et al., 2021). Weppa Farm is a major rice plantation in Nigeria and faces challenges with understanding bird composition in rice plantations and subsequently bird control. There is a need to study the factors influencing avian composition on the rice plantations and make recommendations on biodiversity-friendly agricultural practices in rice production to maximize production and preserve maximum avian biodiversity composition. The aim of this study was to investigate the compositions of bird assemblages in Weppa Farm rice plantation (pre-harvest, during harvest and post-harvest) and the influence of surrounding vegetation on bird abundance. The objectives of this research were;

1. To assess the composition of birds in rice plantations based on farm status

2. Examine the abundance and diversity of different bird-feeding guilds.

3. Investigate how vegetation types surrounding the rice plantation influence the abundance of birds.

## METHODOLOGY

### Study area

Weppa Farm is a 13,000-hectare model mixed farm located near Agenebode, Etsako East local government area in Edo State, Nigeria and was acquired by the Leventis Foundation in 1999 (Leventis foundation, 2021). The farm currently consists of 4,000 hectares of cleared land, an area of conserved woodland and previously uncleared land, which is now being cleared for cultivation. It is a mixed farm with plantations, arable and livestock operations together with integrated industrial processing operation (Leventis foundation, 2021).

**Figure 1:**
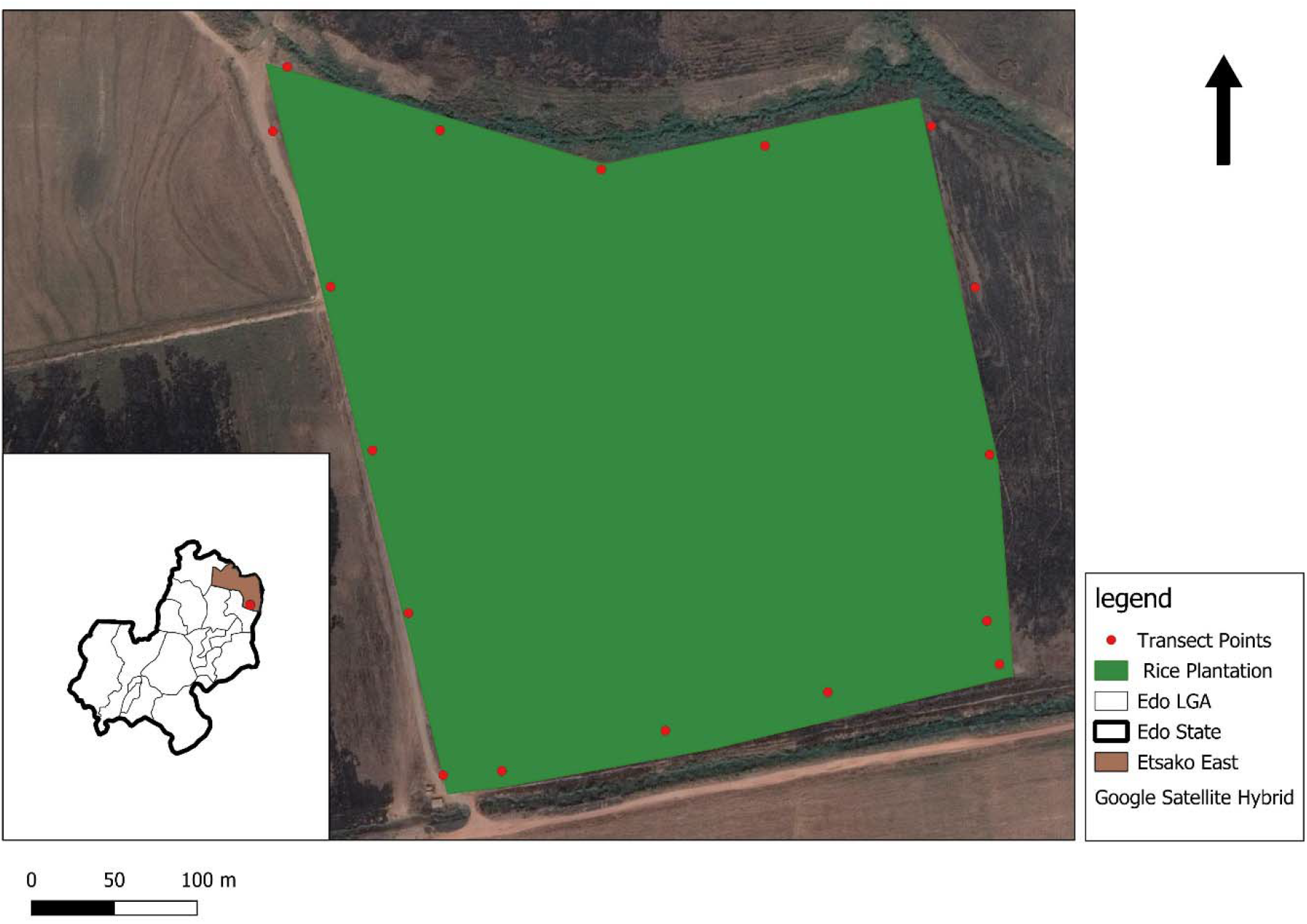
Map of the Edo State showing Rice plantation in Weppa Farms and its environs

### Bird survey

Four line transects were surveyed around the rice plantation (Bibby, 2000). Transects were 400m long and were split into 4, 100m sections (total farm size 139704 sq m). Surveys were done on the rice field before harvest (Unharvested), during harvest (partially harvested) and post-harvest (harvested). Surveys were done in the morning (7:00-9:00 GMT) and repeated in the evening (16:00-18:00 GMT). Vegetation parameters were recorded at the beginning of each section of the transect. The parameters recorded include; surrounding vegetation close to the rice plantation, proportion of weeds to rice on farm, number of trees in adjacent vegetations, number of shrubs in adjacent vegetations and ground cover (%).

### Data analysis

Data was entered on Microsoft excel and analyzed using R statistical software (version 4.3.1). Shannon-Weiner Diversity Index, species abundance, and richness were calculated for each farm status (before, during and after harvest). There were several encountered outliers during the survey. Therefore, in other to deal with issues of model assumptions, a non-parametric (Kruskal-Wallis) test was used to compare bird abundance by farm status, number of trees, feeding guilds and surrounding vegetation.

## RESULTS

A total of 48 different bird species from 6 feeding guilds were recorded. Granivorous birds were the most abundant feeding guild (n=599). The Unharvested rice plantation had the highest species abundance (n=1453), highest richness (r=37) but not diversity. The harvested farm had the highest diversity (H=5.33) though not the highest richness and abundance. The surrounding vegetations had a significant effect on bird abundance.

**Figure 2:**
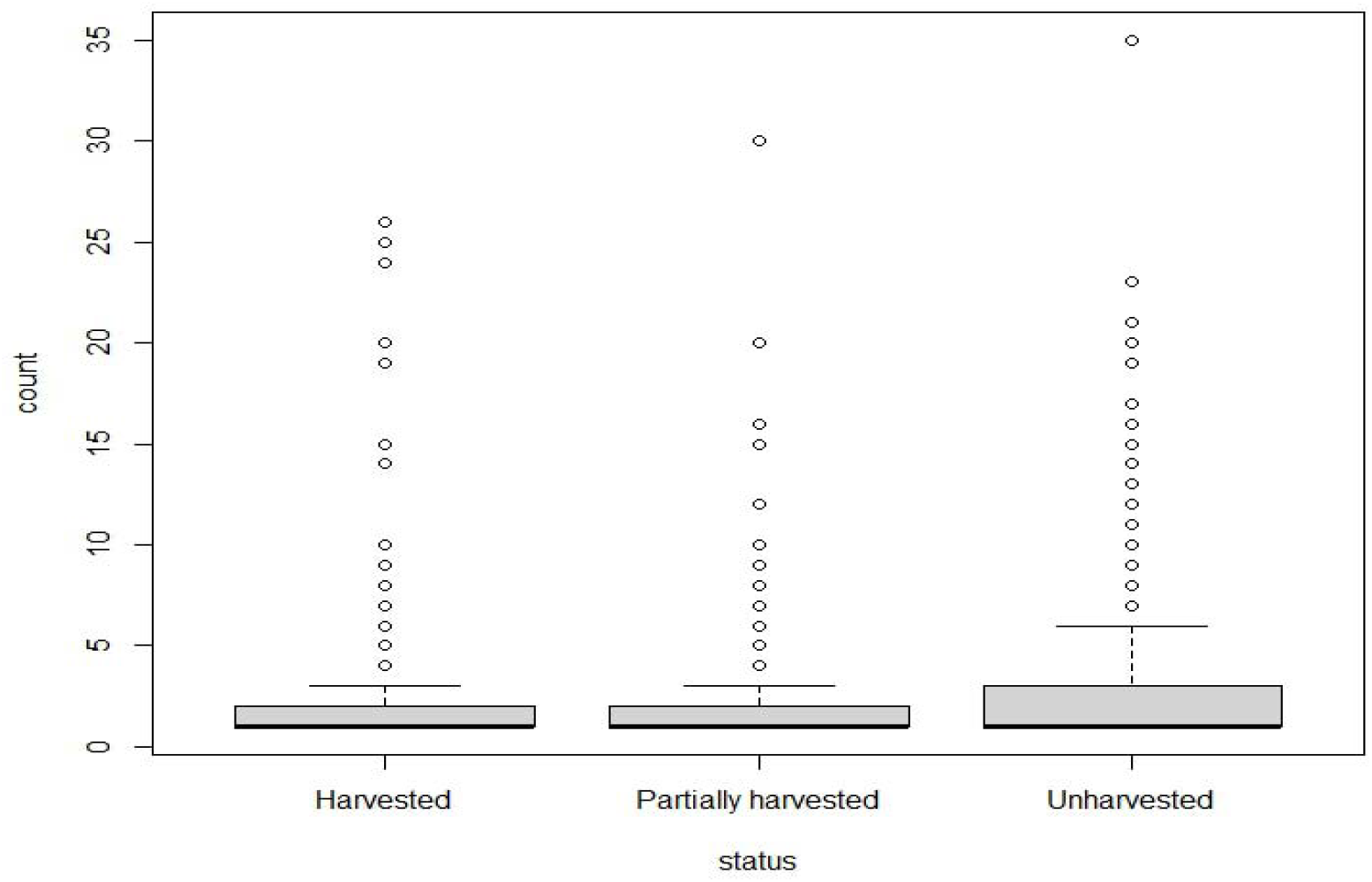
Bird total abundance in the different farm statuses

**Table 1:**
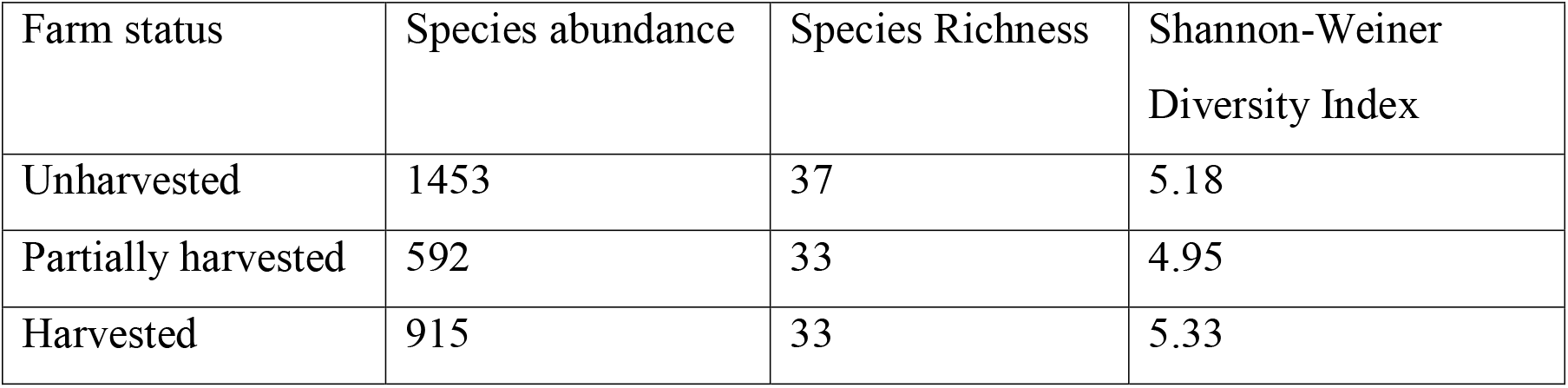
Species abundance, richness and diversity in the farm statuses.

**Figure 3:**
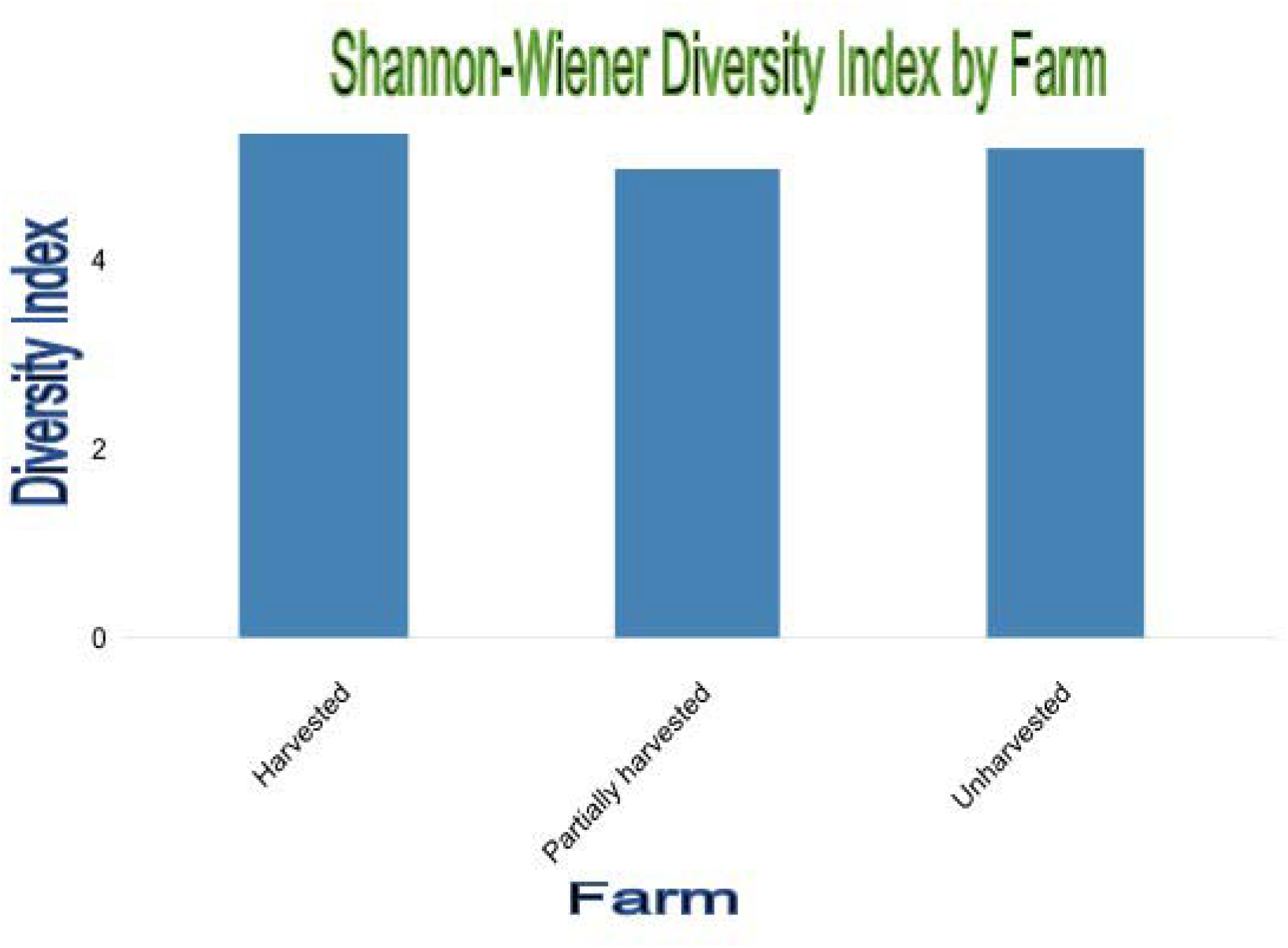
Shannon-Weiner diversity index in the farm statuses

**Table 2:** Effects of surrounding conditions on bird abundance.

| Variable | df | H | P value |
| --- | --- | --- | --- |
| Feeding guild | 5 | 107.89 | $<0.001$ |
| Surrounding vegetation | 4 | 41.202 | $<0.001$ |
| Number of trees | 2 | 12.251 | 0.00219 |
| Number of shrubs | 5 | 47.78 | $<0.001$ |
| Farm status | 2 | 7.8347 | 0.01989 |

H=Kruskal-Wallis chi-squared index

There was a significant difference (*<0*.*001*) in the feeding guilds of the birds recorded in the three farm statuses. Granivores were the most abundant while Nectarivores were the least. Number of trees and number of shrubs caused a significant increase in bird abundance both had significant effect on bird abundance, p=0.00219 and p*<0*.*001* respectively. Adjacent vegetations with a greater number of trees had more avian abundance. Avian abundance significantly differed by farm status (p=0.01989). The unharvested had the highest abundance while the partially harvested had the least.

**Figure 4:**
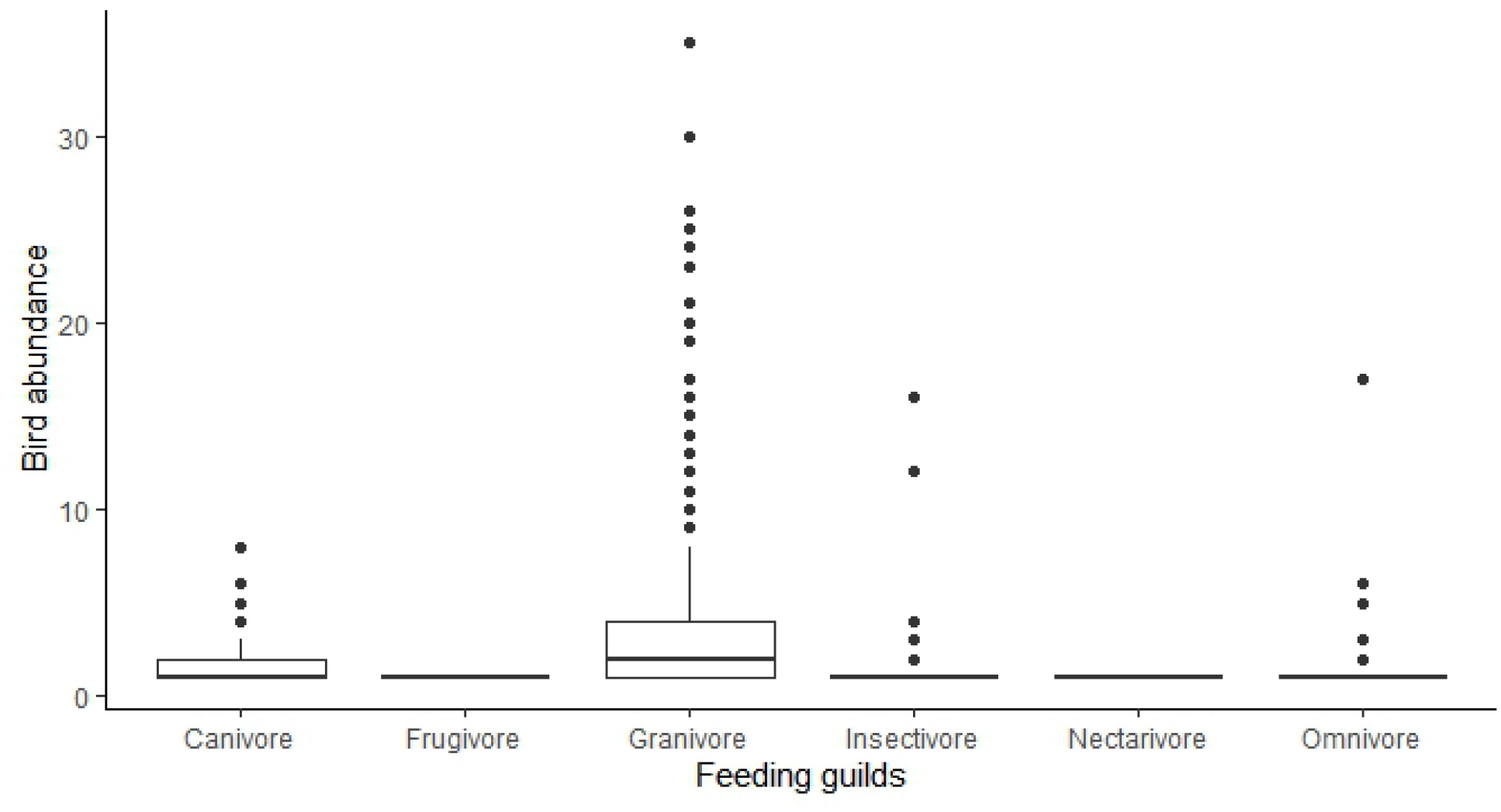
Bird abundance by feeding guild.

Granivorous birds were the most abundant species mainly dominated by weavers followed by insectivores. Nectarivores and frugivores had just 1 and 2 individuals respectively.

**Table 3:** Bird abundance by feeding guilds in the three farm statuses.

| Farm status | Guild | Abundance |
| --- | --- | --- |
| Harvested | Carnivore | 128 |
|  | Granivore | 689 |
|  | Insectivore | 43 |
|  | Omnivore | 55 |
| Partially harvested | Carnivore | 29 |
|  | Granivore | 467 |
|  | Insectivore | 65 |
|  | Nectarivore | 1 |
|  | Omnivore | 30 |
| Unharvested | Carnivore | 38 |
|  | Frugivore | 2 |
|  | Granivore | 1275 |
|  | Insectivore | 82 |
|  | Omnivore | 56 |

**Figure 5:**
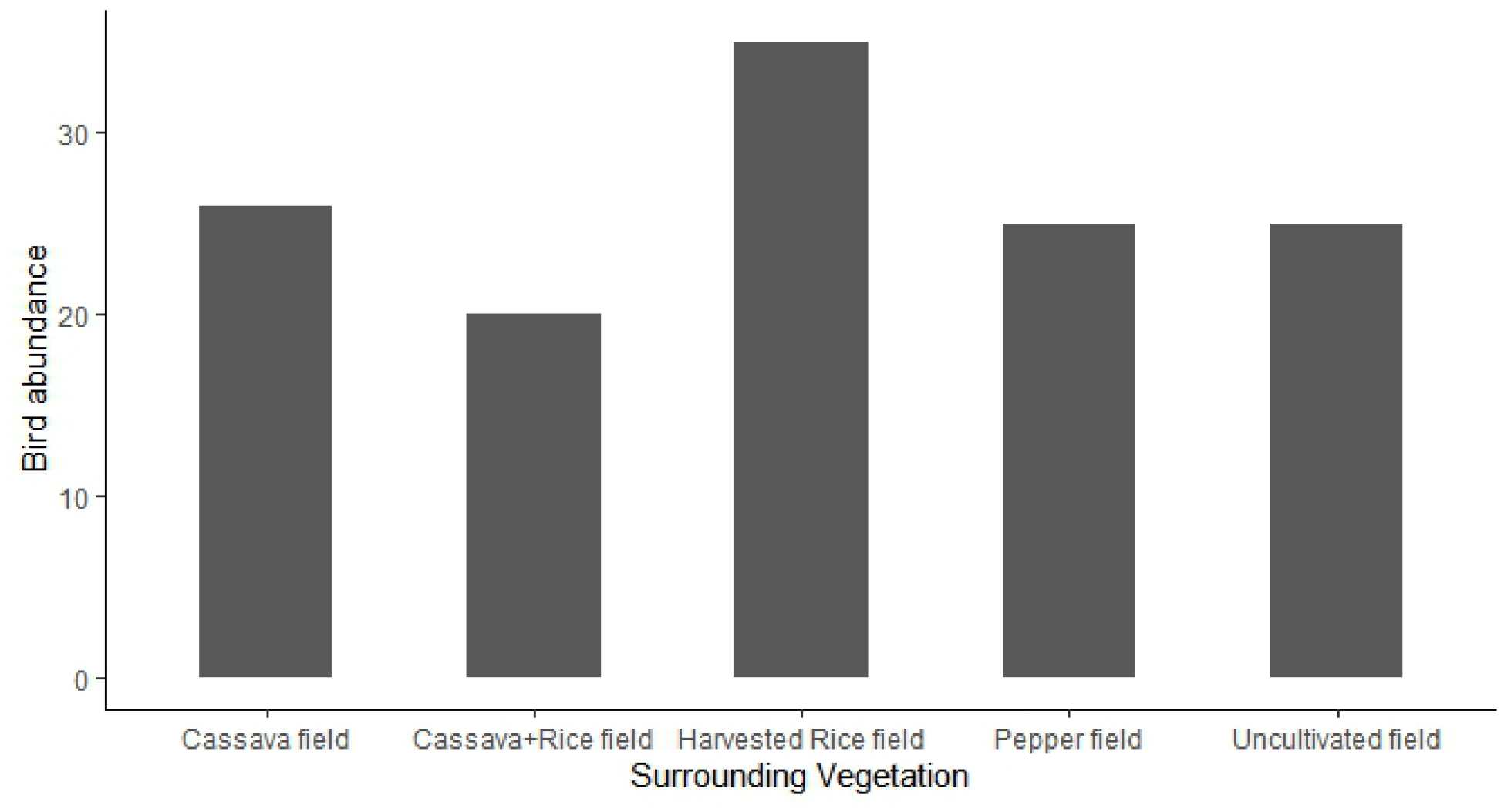
Bird abundance by different surrounding vegetations.

**Figure 6:**
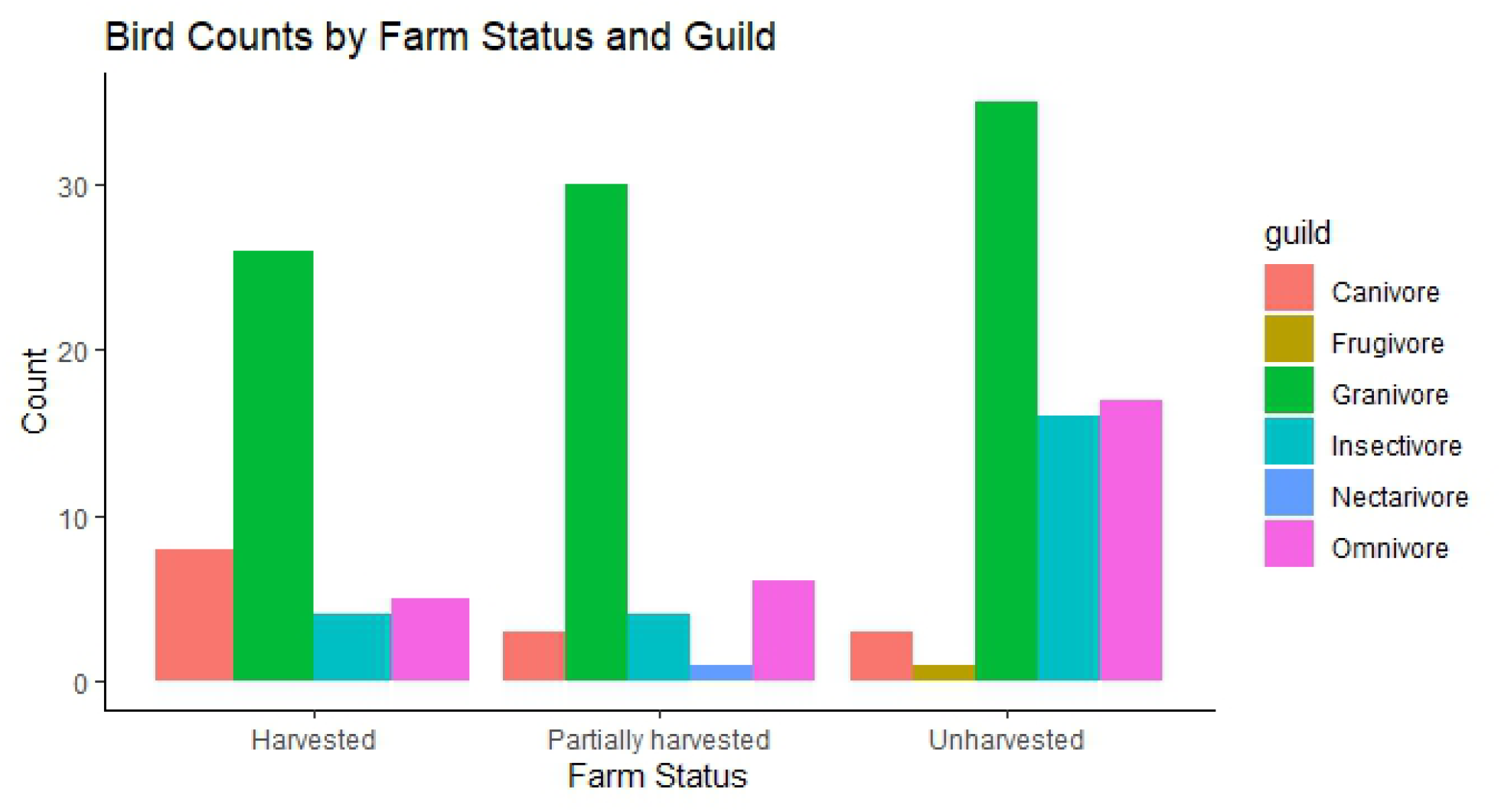
Comparison of feeding guild abundance in harvested, partially harvested, and unharvested farm.

**Figure 7:**
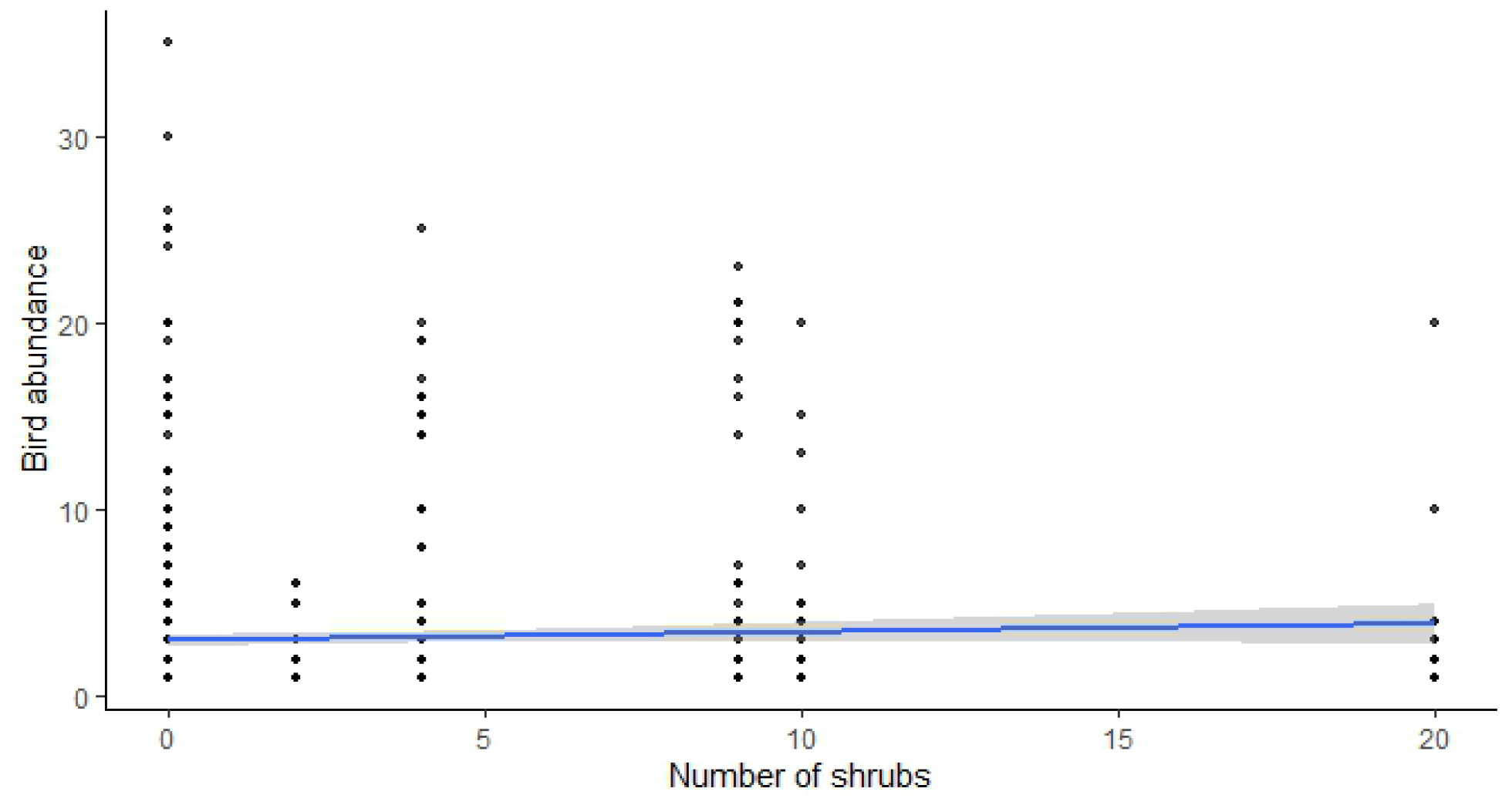
Relationship between the number of shrubs and bird abundance

**Figure 8:**
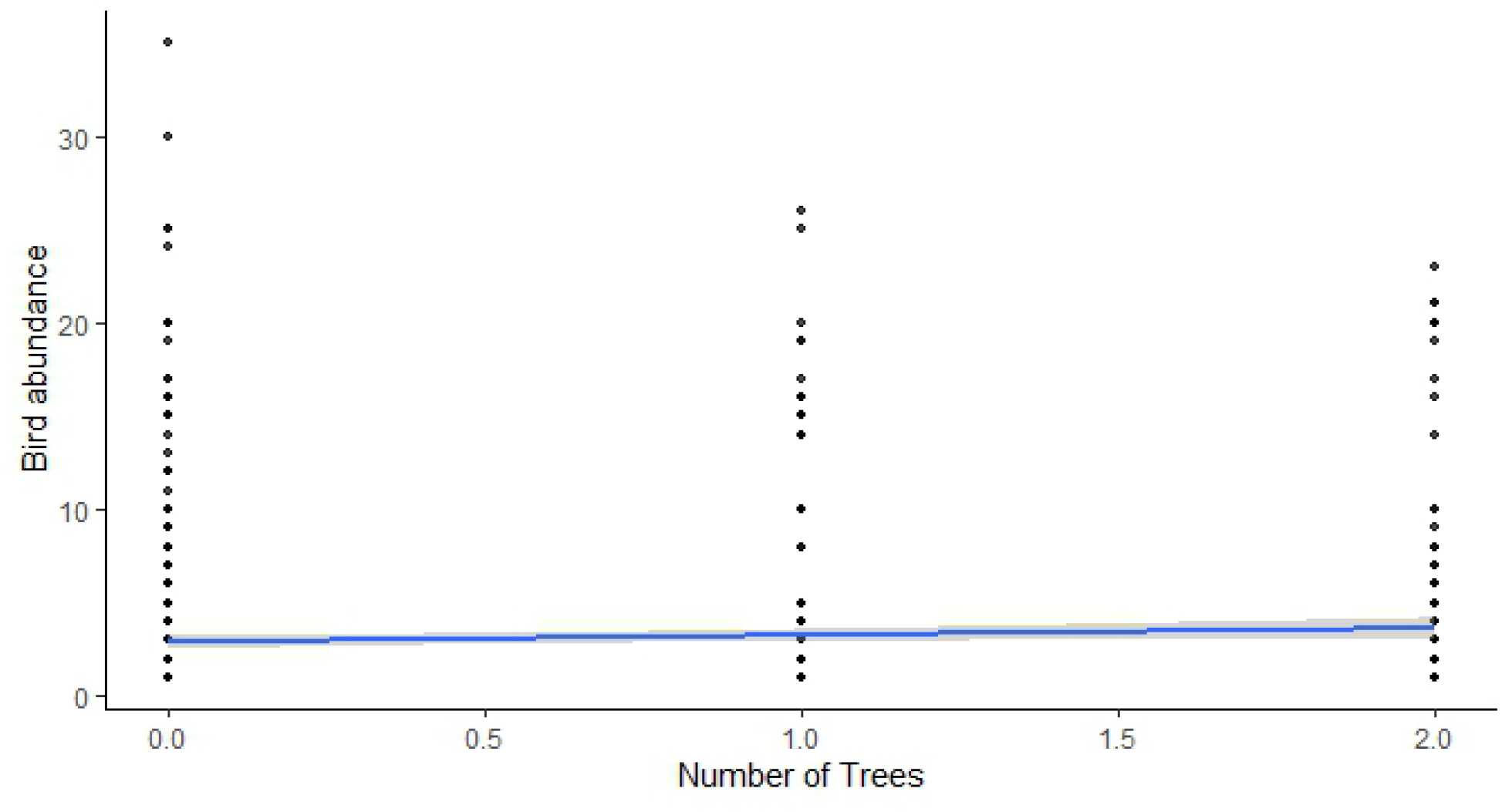
Relationship between number of trees and bird abundance.

## DISCUSSIONS

Species abundance was highest in the Unharvested rice plantation and least in the harvested farms. The unharvested farm had the highest species richness while the harvested farm had the least species richness. The highest species diversity was recorded in the harvested farm while the partially harvested farm had the least diversity index. The unharvested farm had had abundance of due to large flocks of seed and grain eaters mainly village and (*Ploceus cucullatus*) black headed weavers (*Ploceus melanocephalus*). This conforms with the observations of Amzah et al., (2021) in aerobic rice fields in Malaysia and slight strays away from the observations of (Platt et al., 2021) traditional rice ecosystems in Myanmar where insectivores were the most common feeding guild. The partially harvested and the harvested field had reduced granivore abundance compared to the unharvested fields but highest abundance to other guilds in each field. This could be associated to the fact that there were still grain left overs on the field. Carnivores (n=195) and insectivores (n=190) were the next in abundance. The harvested farm supported attracted high carnivore populations such as Black-headed heron, Hamerkop etc. relative to the other fields. This is due to the activity of the harvested which created spaces proceeded by rainfall increasing the populations of small amphibians and vertebrates. The harvested field had the highest abundance of carnivores (n=128) compared to the Unharvested (38) and partially harvested fields (29). This explains why the harvested field had the diversity index (H=5.33). Post-harvest and seedling stage rice fields when flooded, attract a high number of carnivorous birds, especially from the family *Ardeidae* (Taib et al., 2018). The harvested field had more water-logged areas which supported the growth of small amphibians and other aquatic invertebrates attracting carnivorous birds. These birds prey on aquatic organisms present in the flooded fields (Fujioka et al., 2010). This is in line the observations of (Taib et al., 2018) in east Peninsular Malaysia where Carnivorous birds were the most common in harvested rice plantations. Carnivorous birds particularly benefit from rice fields during the sowing stage, while granivorous waterbirds prefer post-harvest flooded fields (Acosta et al., 2010). Insectivorous birds were most abundant in the unharvested fields (n=82) compared to the Partially harvested (n=65) and the Harvested (n=42) and were the next most abundant feeding guild after granivores. Insectivorous birds have been showed to be the most abundant birds in specific rice plantations. (Platt et al., (2021) case study in Myanmar documented 85 bird species in a traditional rice field, with insectivores being the most common feeding guild. The compositions and abundance of insectivorous birds in rice fields are due to factors such water level, flooding period, rice plant structure, and pesticide (Ibáñez et al., 2010). Insectivorous birds always occupy a significant role in rice ecosystems, providing valuable ecological services serving as biocontrol agents for invading arthropods (Sidhu et al., 2024). The type of vegetation adjacent to the rice field had significant influence on bird abundance (p*<0*.*001*). Adjacent vegetations with most trees and shrubs had high bird abundance.

## CONCLUSIONS

Granivores mainly village weavers were the most abundant species. The were mainly observed foraging and using weeds as nesting material. Insectivorous birds were the second must abundant species and could serve as a biocontrol for arthropod invasions on the farm. Surrounding vegetation influences the abundance of birds in a rice plantation. There was more bird abundance in areas that had trees and shrubs nearby. Investigate the role of insectivores in arthropod pest control and if this improves yield in rice plantations. Observe the feeding behavior of Village weavers and determine their impact on rice yields. Assess the effectiveness of using bird scarers and different acoustic methods in rice fields.

## ACKNOWLEDGMENTS

This work was supported by the management of Weppa Farm and research associates for the A.P Leventis Ornithological Research Institute.

## FUNDING

No funding was received from any source to support this research.

## CONFLICTST OF INTEREST

The authors declare no conflicts of interest.

## AUTHOR CONTRIBUTION

I.I, B.A and D.T were involved in designing the research and data collection and manuscript write up. I.I was involved in data analysis. All authors were involved in manuscript review and consented on the publication.

## Notes

### Competing Interest Statement

The authors have declared no competing interest.

## REFERENCES

Acosta, M., Mugica, L., Blanco, D., López-Lanús, B., Dias, R. A., Doodnath, L. W., & Hurtado, J. (2010). Birds of Rice Fields in the Americas. Waterbirds, 33(sp1), 105–122. 10.1675/063.033.s108

Adjah, K. L., Asante, M. D., Toure, A., Aziadekey, M., Amoako-Andoh, F. O., Frei, M., Diallo, Y., & Agboka, K. (2022). Improvement of Rice Production under Drought Conditions in West Africa: Application of QTLs in Breeding for Drought Resistance. Rice Science, 29(6), 512–521. 10.1016/j.rsci.2022.06.002

Amira, N., Rinalfi, T., & Azhar, B. (2018). Effects of intensive rice production practices on avian biodiversity in Southeast Asian managed wetlands. Wetlands Ecology and Management, 26(5), 865–877. 10.1007/s11273-018-9614-y

Amzah, B., Baki, R., & Yahya, M. H. (2021). AVIAN SPECIES COMPOSITION PROFILE AND FEEDING GUILDS UNDER THE AEROBIC RICE FIELD. Jurnal Hama Dan Penyakit Tumbuhan Tropika, 21(1), 63–71. 10.23960/j.hptt.12163-71

Bibby, C. J., B. N. D., H. D. A., and M. S. H. (2000). Bird Census Techniques (2nd ed). Academic Press.

Fujioka, M., Don Lee, S., & Kurechi, M. (2010). Bird use of Rice Fields in Korea and Japan. Waterbirds, 33(sp1), 8. 10.1675/063.033.s102

Ibáñez, C., Curcó, A., Riera, X., Ripoll, I. I., & Sánchez, C. (2010). Influence on Birds of Rice Field Management Practices during the Growing Season: A Review and an Experiment. https://api.semanticscholar.org/CorpusID:86521461

Leventis foundation. (n.d.). https://www.leventisfoundation.org/affiliated-foundations-nigeria.

Odoukpe, S. G., Yaokokore Beibro, H. K., Kouadio, P. K., & Konan, M. E. (2014). Dynamique du peuplement des Oiseaux d’une riziculture et ses environs dans la zone humide d’importance internationale de Grand-Bassam. Journal of Applied Biosciences, 79(1), 6909. 10.4314/jab.v79i1.6

Pierluissi, S. (2010). Breeding Waterbirds in Rice Fields: A Global Review. https://api.semanticscholar.org/CorpusID:85785474

Platt, S. G., Win, M. M., Lin, N., Aung, S. H. N., John, A., & Rainwater, T. (2021). Avian species richness in traditional rice ecosystems: a case study from upper Myanmar. Journal of Threatened Taxa, 13(7), 18719–18737. 10.11609/jott.6992.13.7.18719-18737

Sidhu, S. K., Sekhon, G. S., Kumar, S., Kler, T. K., & Chandi, A. K. (2024). Diversity of Insectivorous Avian Species and their Foraging Activities at Ponds in Agricultural Habitats in Punjab, India. Pakistan Journal of Zoology, 56(2). 10.17582/journal.pjz/20220211020232

Sottomayor, M., Palmeirim, A. F., Meyer, C. F. J., de Lima, R. F., Rocha, R., & Rainho, A. (2024). Nature-based solutions to increase rice yield: An experimental assessment of the role of birds and bats as agricultural pest suppressors in West Africa. Agriculture, Ecosystems & Environment, 370, 109067. 10.1016/j.agee.2024.109067

Taib, F. S. M., Shafawati, F., Kamaruddin, & Aswat, H. (2018). The rice-growing cycle influences diversity and species assemblages of birds in the paddy field ecosystem in east Peninsular Malaysia. Pertanika Journal of Tropical Agricultural Science, 41, 1669–1683. https://api.semanticscholar.org/CorpusID:208232259

